# Antagonistic coevolution between hosts and sexually transmitted infections

**DOI:** 10.1101/590232

**Authors:** Ben Ashby

## Abstract

Sexually transmitted infections (STIs) are predicted to play an important role in the evolution of host mating strategies, and vice versa, yet our understanding of host-STI coevolution is limited. Here, I present a model of acute STI infection in populations with ephemeral mating dynamics, where hosts evolve their preference for healthy mates and STIs evolve mortality or sterility virulence. Mate choice readily evolves even though ephemeral mating and acute infections reduce the advantages of mate choice compared to previous theory based on serial monogamy and chronic infections. Selection for mate choice constrains both mortality and sterility virulence, leading to optimal strategies in each population, host polymorphism, or fluctuating selection. I show how the mode of virulence, costs associated with mate choice, recovery, and host lifespan impact on host-STI coevolution, with fluctuating selection most likely when hosts have intermediate lifespans, STIs cause sterility and longer infections, and costs of mate choice are not too high. The results reveal new insights into the evolution of mate choice and how coevolution unfolds for different host and STI life-history traits, providing increased support for parasite-mediated sexual selection as a potential driver of host mate choice, and mate choice as a constraint on STI virulence.

## Introduction

Parasite-mediated sexual selection (PMSS) is predicted to lead to the evolution of reproductive strategies that limit the risk of infection from mating (Hamilton and Zuk 1982; Sheldon 1993; Loehle 1997). By preferentially selecting mates based on cues related to disease, organisms should be able to increase their reproductive success, either because they might choose partners possessing genes which confer resistance to disease (the “good genes” hypothesis; Hamilton and Zuk 1982), or simply because they choose mates that are currently uninfected and hence are a low-risk option (the “transmission avoidance hypothesis”; Loehle 1997). Both hypotheses have been the subject of intense empirical research with varying evidence in support of and against PMSS (Borgia 1986; Borgia and Collis 1989; Clayton 1990, 1991; Hamilton and Poulin 1997; Abbot and Dill 2001; Webberley et al. 2002; Balenger and Zuk 2014; Jones et al. 2015). In some cases females have been found to prefer uninfected males – for example, Clayton (1990) found that female Rock Doves (*Columba livia*) prefer males without lice, suggesting support for PMSS – while in other cases females appear unable to distinguish between infected males – for instance, female milkweed leaf beetles (*Labidomera* clivicollis; Abbot and Dill 2001) and two-spot ladybirds (*Adalia bipunctata*; Webberley et al. 2002), do not avoid males with sexually transmitted mites. In parallel, there has been much theoretical interest in understanding the role of parasites, especially sexually transmitted infections (STIs), in the evolution of host mating strategies, and vice versa (Thrall et al. 1997, 2000; Knell 1999; Boots and Knell 2002; Kokko et al. 2002; Ashby and Gupta 2013; McLeod and Day 2014; Ashby and Boots 2015). STIs are of particular interest as they are inherently tightly linked to host reproduction unlike non-STIs and are more likely to have negative effects on host fecundity (Lockhart et al. 1996). This body of theoretical work has generally predicted that STIs can indeed act as a strong force of selection on host mating strategies. While changes in host mating behaviour arising from PMSS will in turn affect STI evolution, forming a coevolutionary feedback, almost all theoretical studies only consider one-sided adaptation of either the host or the STI with very few predictions for host-STI coevolution (Ashby and Boots 2015; Wardlaw and Agrawal 2019).

In what appears to be the only coevolutionary model of host mate choice and STI virulence to date, Ashby and Boots (2015) proposed a pair formation model of serial monogamy with reciprocal adaptations between host mate choice and sterility virulence (reductions in host fecundity). Crucially, by taking a coevolutionary approach, this study showed that selection against STI virulence due to mate choice was unlikely to lead to a complete loss of mate choice as had been predicted in studies of one-sided adaptation (Knell 1999). By assuming that hosts are able to preferentially choose mates based on visible signs of disease and that more transmissible/virulent STIs are easier to detect, Ashby and Boots (2015) showed that the evolution of mate choice can prevent runaway selection for parasitic castration, leading to either stable levels of choosiness and virulence, or coevolutionary cycling in these traits. These results represent an important first step in understanding the coevolutionary feedbacks that shape host and STI coevolution, with recent work by Wardlaw and Agrawal (2018) exploring similar ideas to consider the effects of host-STI coevolution on sexual conflict.

Previous theory on the coevolution of host mate choice and STI virulence focused on hosts with serially monogamous mating systems that experience chronic, fertility-reducing STIs (Ashby and Boots 2015). These are reasonable assumptions for many host species and STIs: for example, approximately 90% of bird species are thought to be monogamous (Kleiman 1977), and STIs often cause chronic infections, are more likely to cause reductions in fecundity, and typically have less of an impact on mortality than non-STIs (Lockhart et al. 1996; Knell and Webberley 2004). Yet many species do not form monogamous partnerships – for instance, less than 5% of mammals are thought to be monogamous (Kleiman 1977) – and instead have polygamous mating systems which may involve more ephemeral sexual contacts (Ashby and Gupta 2013). Likewise many STIs are known to increase mortality – for example, HIV and syphilis in humans, and dourine in equines (Gizaw et al. 2017) – and to cause acute rather than chronic infections (e.g. *Chlamydia trachomatis* can be cleared by a number of mammalian species; Miyairi et al. 2010). Broadening our understanding of host-STI coevolution therefore requires the development of theory that captures alternative host mating systems and disease outcomes, as it is currently unclear how host mate choice and STI virulence are likely to coevolve when hosts are not serially monogamous and STIs cause acute infections and/or increased mortality.

Of particular interest is the likelihood that moving from serial monogamy to a mating system with ephemeral sexual contacts will weaken selection for mate choice, as the turnover of sexual partners is much higher and so the costs and benefits of choosing any particular partner is expected to be less extreme than in the case of serial monogamy. This is because forming exclusive sexual partnerships can have both positive and negative effects on reproductive success: pairing with an uninfected partner insulates an individual from contracting infection from other members of the population, but pairing with an infected partner will greatly increase the risk of contracting infection and will potentially lower the chance of producing offspring. Thus, the costs of choosing a ‘bad’ (infected) partner under serially monogamous mating will likely be greater than when sexual contacts are ephemeral, and so we might expect selection for mate choice to be especially strong under serial monogamy.

Here, I analyse a simple model of host-STI coevolution under ephemeral mating dynamics with mortality or sterility virulence, and recovery from infection. Using evolutionary invasion analysis, I show how mortality and sterility virulence are constrained by mate choice and when mate choice is likely to evolve under different disease characteristics. I then show that coevolutionary cycling is typically more common under sterility virulence, whereas mortality virulence tends to lead to more stable outcomes. I also identify conditions when evolutionary branching in host mate choice is possible, showing that this outcome appears to be highly constrained. Finally, I examine how costs associated with mate choice, the rate of recovery from infection, and the lifespan of the host impact on host-STI coevolution. Overall, these results show that while serial monogamy, sterility virulence, and chronic infections tend to increase selection for mate choice, parasite-mediated sexual selection is likely to occur for a much broader range of host and STI life-history traits.

## Methods

I model the epidemiological dynamics of a sexually transmitted infection (STI) and the mating dynamics of a well-mixed, hermaphroditic host population according to the following system of ordinary differential equations:

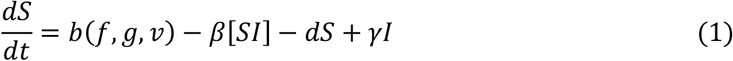

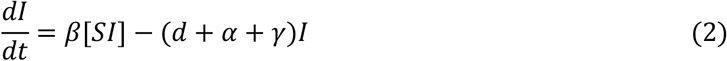

where: ***S*** and ***I*** are the number of susceptible and infected individuals, respectively; *b*(***f***, ***g***, ***𝑣***) is the host birth rate, which depends on the relative fecundity of infected hosts, ***f***, with 0 ≤ ***f*** ≤ 1, the mate choice strategy, ***g*** ≥ 0, and the virulence of the STI, ***𝑣***; ***d*** is the natural mortality rate; ***α*** is the disease-associated mortality rate; ***β*** is the transmission rate per sexual contact, ***γ*** is the recovery rate; and [***XY***] is the mating rate between individuals in classes ***X*** ∈ {***S***, ***I***} and ***Y*** ∈ {***S***, ***I***}. Hereafter, I assume that mortality virulence and sterility virulence may be functions of transmissibility (i.e. ***α*** = ***α***(***β***), ***f*** = ***f***(***β***), ***𝑣*** = ***𝑣***(***β***)) as parasites typically need to damage their hosts or use host resources to produce transmission stages, and the greater the transmissibility of the parasite there is likely to be greater damage caused to the host.

Sexual contacts are assumed to be ephemeral, and the mating dynamics occur as follows. The baseline per-capita mating rate is ***p***, which is independent of the population size, ***N*** = ***S*** + ***I***. This means that larger populations do not have a higher per-capita mating rate than smaller populations. Before deciding whether or not to mate, each host inspects its prospective partner for signs of infection (e.g. through visual or olfactory clues). If the prospective partner is currently uninfected, the probability that the focal host accepts the mate is ***m***_***S***_(***g***), with 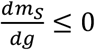. The function ***m***_***S***_(***g***) allows for the fact that hosts may imperfectly assess the condition of other individuals and those who are choosier (higher ***g***, lower ***m***_***S***_(***g***)) may be generally more cautious in their approach to mating, potentially declining healthy prospective partners. If the prospective partner is currently infected, the probability of accepting them as a mate, ***m***_*I*_ (***g***, ***𝑣***(***β***)) with 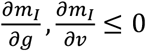, depends on both the choosiness strategy of the host, ***g***, and the virulence, ***𝑣***(***β***), (either sterility ***𝑣***(***β***) **= 1** − ***f***(***β***), or mortality ***𝑣***(***β***) = ***α***(***β***) virulence) of the STI. By causing more damage to their hosts, more virulent STIs may be easier to detect by prospective mates (e.g. due to a general deterioration in health or more visible signs of infection). Note that ***g*** is a dummy variable which I refer to as the mate choice strategy, whereas ***m***_***S***_(***g***) and ***m***_***I***_(***g***, ***𝑣***(***β***)) are the probabilities of accepting an uninfected or infected prospective partner as a mate, respectively. I use a dummy variable for the mate choice strategy so that the host responses to susceptible and infected individuals are correlated. Throughout, it is assumed that the probability of accepting an uninfected individual as a mating partner is at least as large as the probability of accepting an infected individual: ***m***_***S***_(***g***) ≥ ***m***_***I***_(***g***, ***𝑣***(***β***)). Without this assumption, there would never be any advantages to mate choice, as choosier individuals would mate with infected members of the population at a higher rate than less choosy individuals. For the results section I therefore set 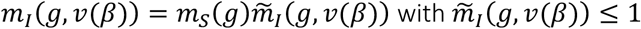 the mate choice response specific to prospective partners who are infected. The mating rates for each combination of the ***S*** and ***I*** classes are given by:

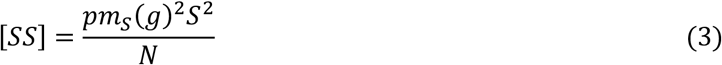

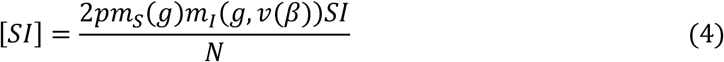

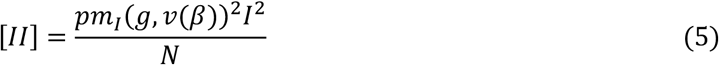

The factor of 2 in the equation for [***SI***] appears when there is mating between individuals in different classes and is required to balance the total mating rate, ***M***:

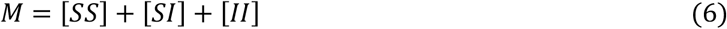

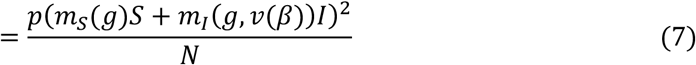

Note that for the specific case when ***m***_***S***_(***g***) = ***m***_***I***_(***g***, ***𝑣***(***β***)) **= 1** (i.e. there is no mate choice), the total mating rate reduces to ***M*** = ***pN***. For the general case, hosts that have mated produce offspring at a total rate of:

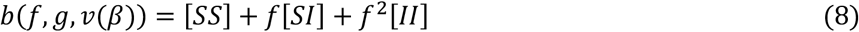

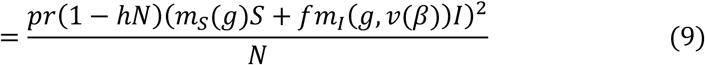

where ***r*** is the maximum reproduction rate per pair and the birth rate is subject to density-dependent competition given by the parameter ***h***.

The disease-free equilibrium (***S***, ***I***) = (***S***^∗^, 0) of this system occurs at:

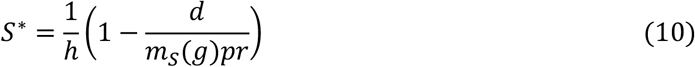

and is viable provided ***m***_***S***_(***g***)***pr*** > ***d*** (i.e. the birth rate is higher than the death rate). A newly introduced STI will spread in a susceptible population when the basic reproductive ratio, ***R***_0_(***g***, ***β***) is greater than 1, where:

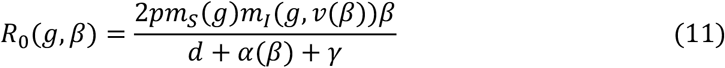

The above model describes the dynamics when there is only one host type and one STI type in the population. In polymorphic populations, the dynamics for ***n***_***h***_ hosts each with trait ***g***_***i***_, where ***i*** ∈ {1, …, ***n***_***h***_} and ***n***_***p***_ STIs each with transmissibility ***β***_***j***_ and virulence ***𝑣***_***j***_ where ***j*** ∈ {1, …, ***n***_***p***_}, are fully described by the following system of ordinary differential equations:

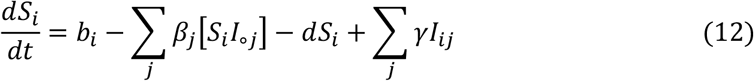

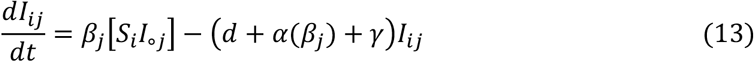

where 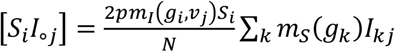 is the total mating rate between susceptible hosts with trait ***g***_***i***_ and all hosts infected by STIs with traits ***β***_***j***_ and ***𝑣***_***j***_, and the birth rate for each host type is:

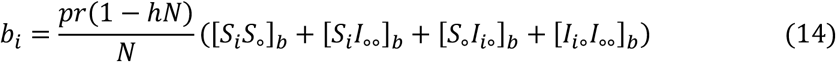

where for notational convenience:

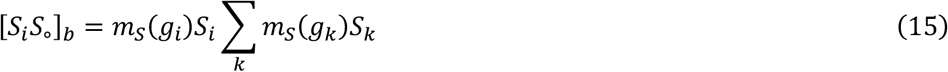

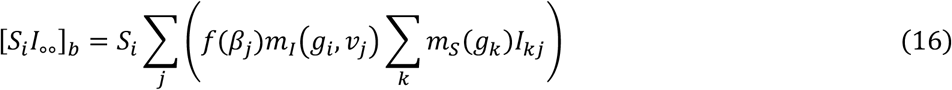

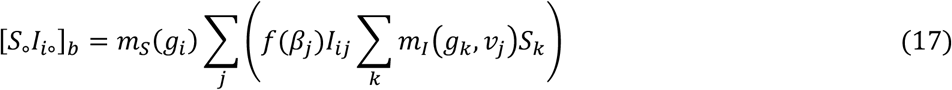

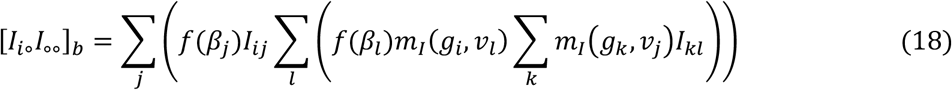

This model is related to the pair formation framework proposed in Ashby and Boots (2015), but there are two key differences. First, the model in Ashby and Boots (2015) assumes there is serial monogamy with separate pools of paired and unpaired individuals. Unpaired individuals form potentially long-term sexual partnerships at certain rates that terminate when either partner decides to seek a new mate or one partner dies, at which point the former partner(s) return to the pool of unpaired individuals. Here, I focus on a single pool of individuals with ephemeral sexual contacts, which is analogous to having infinite pair dissolution rates and instantaneous reproduction. However, it is not possible to move directly from the previous model to the present one by letting the pair dissolution rates tend to infinity, as this would mean individuals are never in the paired state and reproduction only occurs while individuals are paired in Ashby and Boots (2015). The transition to ephemeral mating may have effects on both the costs and benefits of mate choice. For example, when sexual contacts are ephemeral rather than serially monogamous, individuals no longer spend a period of time paired (and hence isolated) from the rest of the population, which means that on average hosts will come into contact with a larger number of mates. Under serial monogamy, choosing a ‘bad’ partner is very costly because of this isolation effect (higher risk of infection due to sustained contact, fewer offspring), but under ephemeral mating the risk of eventually coming into contact with an infected mate will be higher (more rapid partner turnover).

The second major difference is in the nature of the infection: the model in Ashby and Boots (2015) assumes STIs cause reductions in host fecundity rather than impact on mortality, and that individuals are unable to recover once infected. Relaxing these assumptions in the present model, by allowing STIs to cause mortality and hosts to clear infection (as is the case for many STIs; e.g. Miyairi et al. 2010; Gizaw et al. 2017), will reveal how a broader range of STI life-history traits affect coevolution with host mate choice. An additional difference in the current model is that the mate choice functions have been generalised to ***m***_***S***_(***g***) and ***m***_***I***_(***g***, ***𝑣***(***β***)) for easier interpretation and so that alternative mate choice relationships can be readily explored.

For one-sided adaptation (either host or STI evolution) I use evolutionary invasion analysis to determine the long-term trait dynamics (Geritz et al. 1998). This assumes that mutations have small phenotypic effects and are sufficiently rare so that the system has reached a stable state before a new mutant emerges. I solve the dynamics numerically as the system is intractable to classical methods of stability analysis. For coevolution, I relax the assumptions of the evolutionary invasion analysis by having a discrete set of host and STI types and by allowing mutations to occur before the system has reached a stable state in simulations (source code in the *Supporting Information*).

## Results

### Parasite evolution

The invasion fitness of a rare mutant strain of the STI (subscript ***m***) in a population at equilibrium ***N***^∗^ = ***S***^∗^ + ***I***^∗^ is:

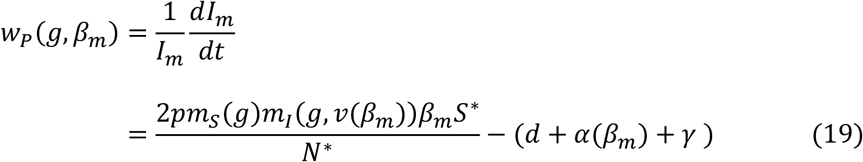

The mutant STI can only invade when ***w***_***p***_(***g***, ***β***_***m***_) > 0, which requires

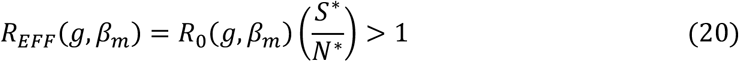

where ***R***_***EFF***_(***g***, ***β***_***m***_) is the effective reproductive ratio. Since STI fitness can be written in this form, we know that parasite evolution maximises ***R***_0_ (Lion and Metz 2018). The STI will evolve in the direction of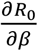 until ***β*** is maximised at 1, one or both populations are driven extinct, or a singular strategy ***β***^∗^ is reached at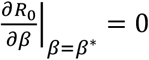, which requires:

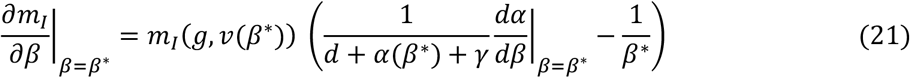

In general, ***m***_***I***_(***g***, ***𝑣***(***β***)) and ***f***(***β***) will be decreasing (or constant) functions of ***β*** and ***α***(***β***) will be an increasing (or constant) function. In the absence of mate choice (***m***_***S***_(***g***) = ***m***_***I***_(***g***, ***𝑣***(***β***)) **= 1**), a *continuously stable strategy* (CSS) – analogous to an *evolutionary stable strategy* or ESS – can only exist when ***α***(***β***) is concave up (i.e. mortality virulence accelerates with the transmission rate). In the presence of mate choice, however, a CSS can exist under a broader set of conditions, such as concave down mortality-transmission trade-offs and with sterility-transmission trade-offs. This is clear from the equation for ***R***_0_(***g***, ***β***) (equation 11), which features the product of ***m***_***I***_(***g***, ***𝑣***(***β***)) and ***β*** (i.e. the product of decreasing and increasing functions of ***β***) (Fig. 1A).

**Figure 1.**
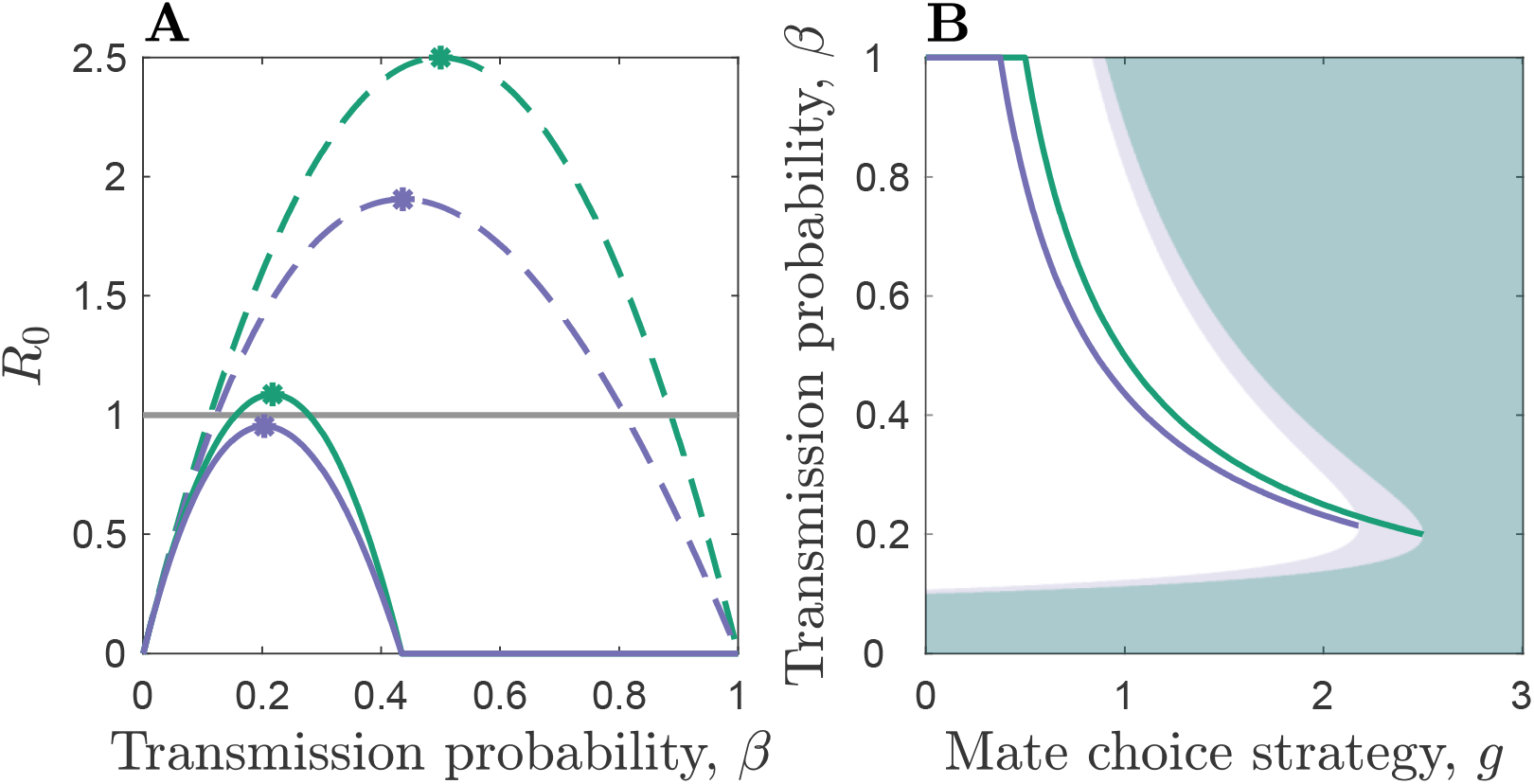
STI evolution in response to host mate choice for sterility (green) and mortality (purple) virulence. (A) Basic reproductive ratio, ***R***_0_, for weaker (dashed; ***g*** **= 1**) and stronger (solid; ***g*** **= 2.3**) mate choice strategies. The horizontal line indicates the extinction threshold for the STI. (B) Curves show the continuously stable strategies (CSSs, i.e. ***β***^∗^) for a given mate choice strategy, ***g***. The corresponding shaded regions show where the STI is unviable. Mate choice and virulence functions as described in the text. Parameters: ***m***_***S***_(***g***) **= 1**, ***γ*** **= 0.5**, ***η*** **= 1**, ***κ*** **= 1**, ***d*** **= 1**, ***h*** **= 10^−6^**, ***p*** **= 7.5**, ***r*** **= 20**.

To illustrate the above, suppose first that either the STI does not impact on fertility (***f***(***β***) **= 1**) or that sterility virulence is constant 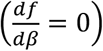 and mortality virulence is a linear function of the transmission probability such that ***𝑣***(***β***) = ***α***(***β***) = ***κβ***. For example, these functions could relate to situations where doubling the number of transmission stages doubles the damage caused to the host (i.e. virulence has an additive or proportional effect on the host). When there is no mate choice the STI will evolve to maximise ***β***. If, however, mate choosiness is a linear function of mortality virulence (i.e. increasing damage has a linear effect on the probability of accepting a mate) such that 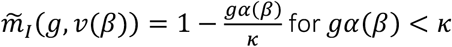 for ***gα***(***β***) < ***κ*** and 0 otherwise, then a singular strategy exists at

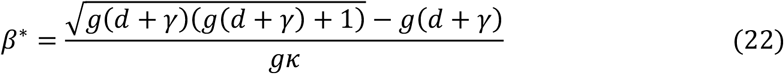

(Fig. 1B) which is always evolutionarily stable since:

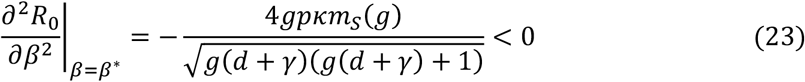

Now suppose instead that mortality virulence is zero (***α***(***β***) **= 0**) or constant 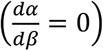 and sterility virulence is a function of the transmission probability such that ***𝑣***(***β***) **= 1** − ***f***(***β***) and ***f***(***β***) **= 1** − ***ηβ*** where 0 < ***η*** ≤ 1 controls the strength of the relationship. If host mate choosiness is a linear function of sterility virulence such that 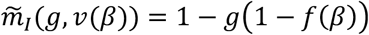 (as with mortality virulence this could correspond to proportionality between the number of transmission stages and the damage caused to the host), then the singular strategy occurs at 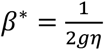 (Fig. 1B), which again is always evolutionarily stable since:

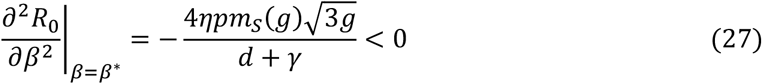

All else being equal, the effects of mate choice on ecology and evolution of the STI under sterility and mortality virulence are qualitatively similar (Fig. 1). However, since mortality virulence causes an additional reduction in ***R***_0_ compared to sterility virulence due to the presence of ***α***(***β***) in the denominator (equation 11), a given level of mate choice will have a greater impact on an STI that causes mortality virulence. This can be seen in Fig. 1, where both ***R***_0_ and the evolved probability of transmission, ***β***^∗^, of and STI are always lower under mortality virulence and the region of viability is smaller compared to STIs that cause sterility virulence.

In summary, even under ephemeral mating and when hosts are able to recover from infection, host mate choice can constrain the evolution of mortality or sterility virulence in an STI, but the effects on STIs that cause mortality virulence will tend to be greater, leading to lower disease prevalence and selection for slightly lower transmissibility for a given level of mate choice.

## Host evolution

The initial dynamics of a rare mutant host (subscript ***m***) in a resident population at equilibrium are given by:

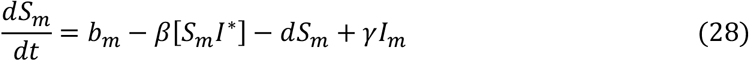

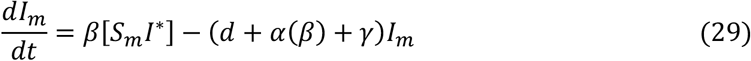

with

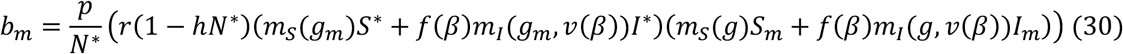

Using the next-generation method (see *Supporting Information;* Hurford et al. 2010), it can be shown that host fitness is sign-equivalent to

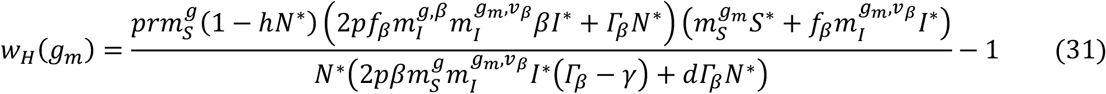

where ***Γ**_β_* = ***d*** + ***α***(***β***) + ***γ***, ***f***_*β*_ = ***f***(***β***), 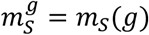 and 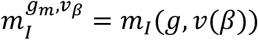 for the sake of brevity. The host will evolve in the direction of 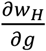 until ***g*** is minimised at 0, one or both populations are driven extinct, or a singular strategy, ***g***^∗^, is reached at 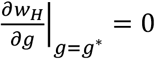.

Suppose initially that mate choice only affects prospective partners that are infected (***m***_***S***_(***g***) **= 1**) and that mate choice of infected individuals is a linear function of virulence, ***𝑣***(***β***), such that 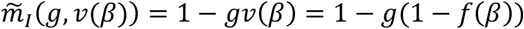 for sterility virulence and 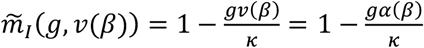 for mortality virulence. This means that the probability of accepting an infected mate is proportional to the damage caused by the STI. In this scenario there may be one or two singular strategies. The singular strategy at 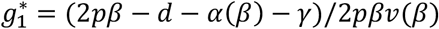 always exists, and corresponds to the point where the host drives the STI extinct. The second singular strategy 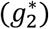, if it exists, is an evolutionary repeller with 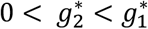, in which case the outcome depends on the initial conditions, with 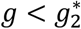 causing selection against mate choice, and 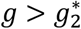 leading to STI extinction due to mate choice (Fig. 2A-B).

**Figure 2.**
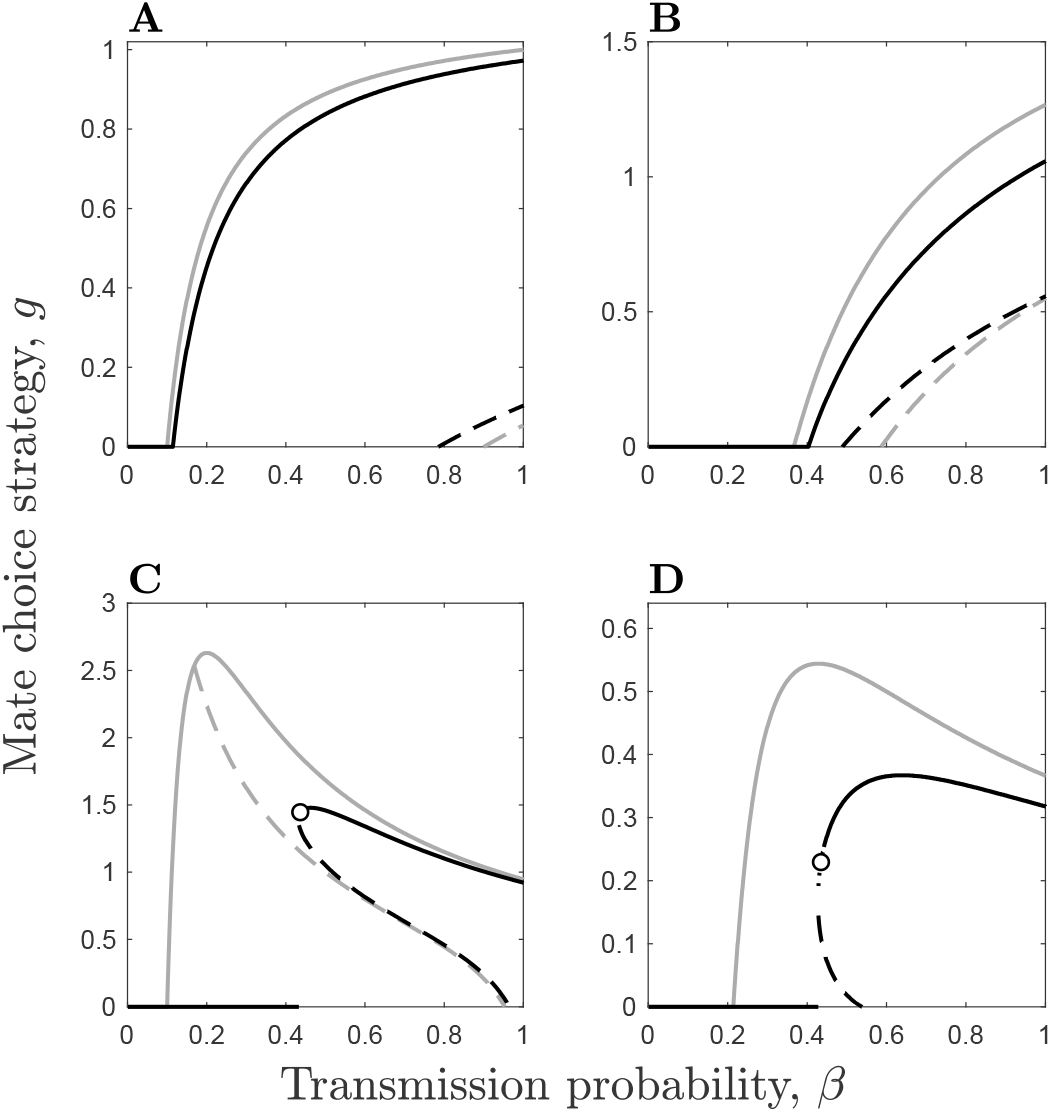
Evolution of the host mate choice strategy, ***g***, in the presence of a non-evolving STI. Solid lines correspond to evolutionary attractors, dashed lines to evolutionary repellers, and circles to branching points, in the presence (black; ***ζ*** = 0.1) and absence (grey; ***ζ*** = 0) of host costs, with ***m***_***S***_(***g***) = 1 − ***ζg***. The component of mate choice specific to infected individuals is as described in the text. Only one type of virulence (mortality or sterility) is assumed to occur in each panel with mate choice of infected individuals based on: (A) fixed sterility virulence, ***𝑣***(***β***) = 0.9; (B) fixed mortality virulence, 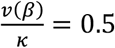; (C) variable sterility virulence, ***𝑣***(***β***) = 1 − ***f***(***β***) = ***βη***; (D) variable mortality ***κ*** virulence, ***𝑣***(***β***) = ***α***(***β***) = ***βκ***. Remaining parameters as described in Fig. 1, except ***κ*** = 8.

Mate choice is likely to evolve for intermediate transmission probabilities when virulence is fixed (Fig. 2A-B). When the probability of transmission is small, the STI is unable to spread even in the absence of mate choice (***R***_0_ < 1). When the probability of transmission is close to 1, there may be selection against weak mate choice caused by the evolutionary repeller. This is because disease prevalence is high and so most attempted matings are with infected individuals, meaning that even weak mate choice dramatically reduces the mating rate for invading host mutants compared to the resident population. If, however, there is already a sufficient level of mate choice in the resident population (i.e. the initial conditions are above the repeller), disease prevalence is sufficiently low to allow runaway selection for mate choice, eventually driving the disease extinct. This pattern is similar regardless of whether virulence has fixed effects on mortality or sterility (Fig. 2A-B). When sterility or mortality virulence is linked to the transmission probability, the dynamics are more complex (Fig. 2C-D). Notably, the threshold for driving the STI extinct is lower at high transmission probabilities because virulence (and hence the effects of mate choice) are also stronger. An evolutionary repeller may exist, but it now occurs for intermediate values of ***β***.

The system is intractable to classical analysis when mate choice also affects prospective partners that are susceptible (i.e. there is a ‘cost’ of being choosy, with ***m***_***S***_(***g***) < 1 for ***g*** > 0), and so one must find the evolutionary dynamics using numerical analysis. While many of the results are qualitatively similar to the no-cost scenario, there are some notable exceptions. In particular, if the host evolves mate choice then it no longer drives the STI extinct, and is instead likely to reach a *continuously stable strategy* (CSS) with the STI endemic in the population. When virulence is a correlated with transmission, the host only evolves mate choice *de novo* at sufficiently high values of ***β*** (Fig. 2C-D). Additionally, there is a very small region of parameter space at intermediate values of ***β*** that can yield evolutionary branching, with stable coexistence between two host types: one which exhibits moderate mate choice and the other which does not discriminate against infected mates.

## COEVOLUTION

I now consider the coevolution of the host and the STI, focusing on how the costs associated with mate choice (***ζ***), the recovery rate (***γ***), and the relative lifespan of the host 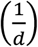 interact with the mode of virulence (sterility or mortality) and the shape of the mate choice response to infected individuals. The model exhibits the same range of qualitative outcomes under both sterility and mortality virulence: (i) *co-continuously stable strategies* (co-CSSs) where STI virulence may be constrained by host mate choice at a stable equilibrium (Fig. 3A-B); (ii) coevolutionary cycling, whereby host and STI phenotypes fluctuate through time (Fig. 3C-D); and (iii) a stable level of virulence in the STI coupled with evolutionary branching in the host, where more and less choosy hosts are able to coexist (Fig. 3E-F). Note that the STI did not branch under any conditions.

**Figure 3.**
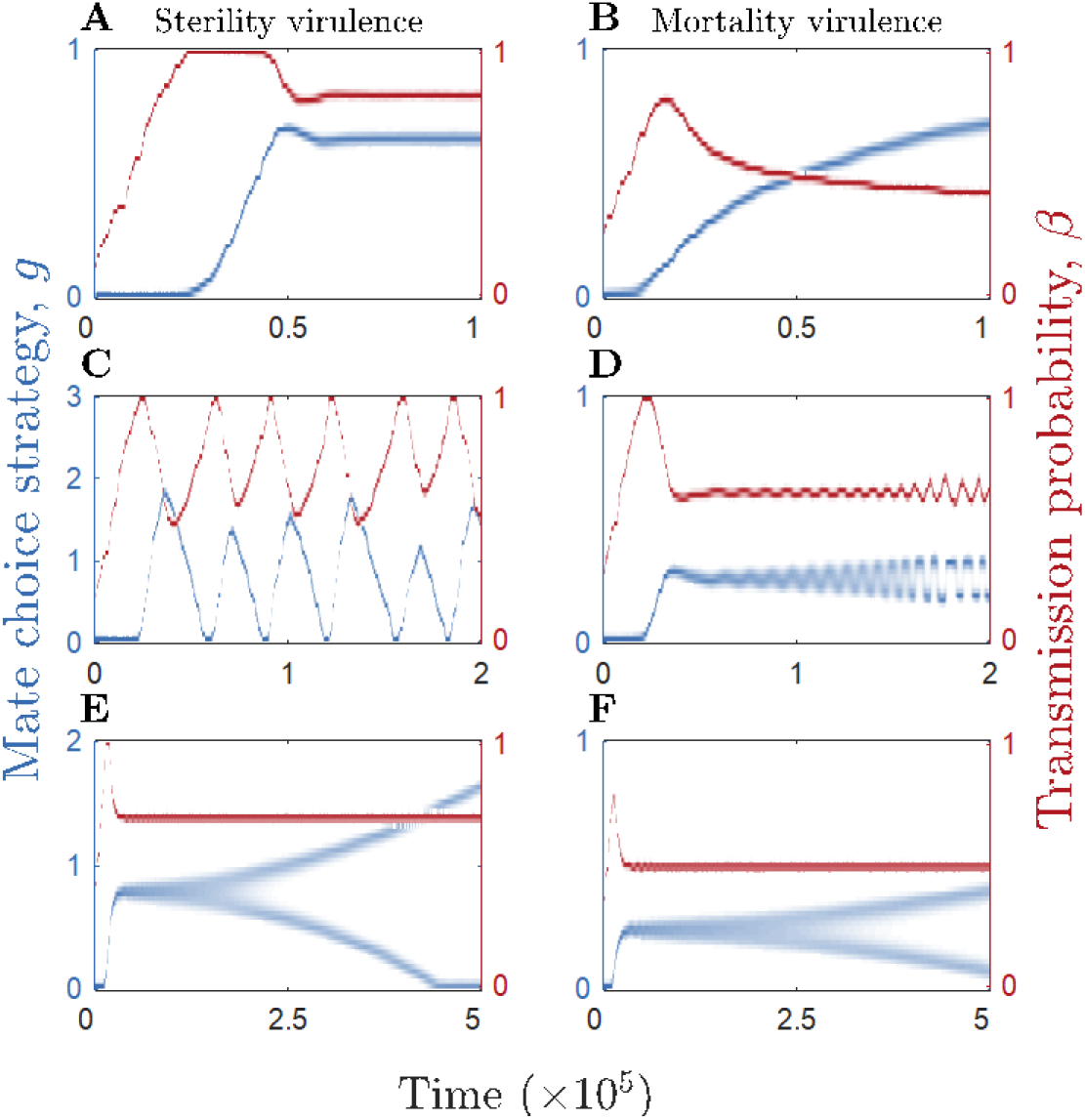
Coevolutionary dynamics between the host mate choice strategy, ***g*** (blue), and the STI transmission probability, ***β*** (red), under sterility (left column, ***𝑣***(***β***) = 1 − ***f***(***β***) = ***βη***) and mortality (right column, ***𝑣***(***β***) = ***α***(***β***) = ***βκ***) virulence. (A-B) The host and STI evolve to co-continuously stable strategies (co-CSS). (C-D) The host and STI exhibit coevolutionary cycling (fluctuating selection). (E-F) The STI evolves to a continuously stable strategy (CSS) but the host branches into two strategies, one with high mate choice and the other with low/no mate choice. Fixed parameters: ***η*** = 0.95, ***κ*** = 8, ***d*** = 1, ***h*** = 10^−6^, ***p*** = 7.5, ***r*** = 20. Other parameters: (A) ***γ*** = 0.5, ***ζ*** = 0.5. (B) ***γ*** = 0.3, ***ζ*** = 0.03. (C) ***γ*** = 0.5, ***ζ*** = 0.1. (D) ***γ*** = 0.25, ***ζ*** = 0.15. (E) ***γ*** = 3, ***ζ*** = 0.1. (F) ***γ*** = 0.3, ***ζ*** = 0.075. Mating functions: (A) 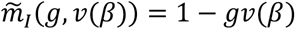. (C, E) 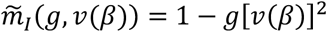. (B, D, F) 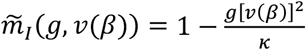.

The coevolutionary dynamics under sterility and mortality virulence are summarised in Fig. 4–5 and Table 1. Overall, higher costs tend to suppress mate choice and allow higher levels of virulence to evolve (Fig. 4A, 5A) and faster recovery rates have a stabilising effect on the dynamics (Fig. 4B, 5B), as do both short and long host lifespans (Fig. 4C, 5C). Although both sterility and mortality virulence produce the same range of qualitative coevolutionary outcomes, there are some notable differences between the two scenarios. First, coevolutionary cycling is much more common under sterility virulence than mortality virulence, with the latter more likely to lead to stable equilibria instead. Even when mortality virulence does produce coevolutionary cycling, the amplitude of the cycles tends to be smaller compared to sterility virulence (Fig. 3C-D). Second, mate choice requires much lower costs (***ζ***) to evolve when the STI causes mortality virulence (Fig. 4A, 5A). Third, higher costs cause qualitatively different transitions in the coevolutionary dynamics, from cycling to stable strategies in the case of sterility virulence and from stable strategies to dimorphism and cycling in the case of mortality virulence (Fig. 4A, 5Aii). Fourth, while mate choice peaks for intermediate host lifespans in the case of sterility virulence (Fig. 4C), under mortality virulence mate choice generally decreases (or for a narrow window becomes dimorphic) as host lifespan increases (Fig. 5C). These general differences in outcomes are broadly consistent whether mate choice is linearly or non-linearly dependent on virulence (Fig. 4–5). Biologically, the non-linear mate choice functions shown here mean that the effects of mate choice accelerate with virulence. There are some differences in outcomes between the linear and non-linear versions, however. For example, when greater virulence is associated with an acceleration in mate choice there is usually a greater potential for coevolutionary cycling under sterility virulence (Fig. 4 top vs bottom row) and for coevolutionary cycling and evolutionary branching under mortality virulence (Fig. 5 top vs bottom row).

**Table 1.**
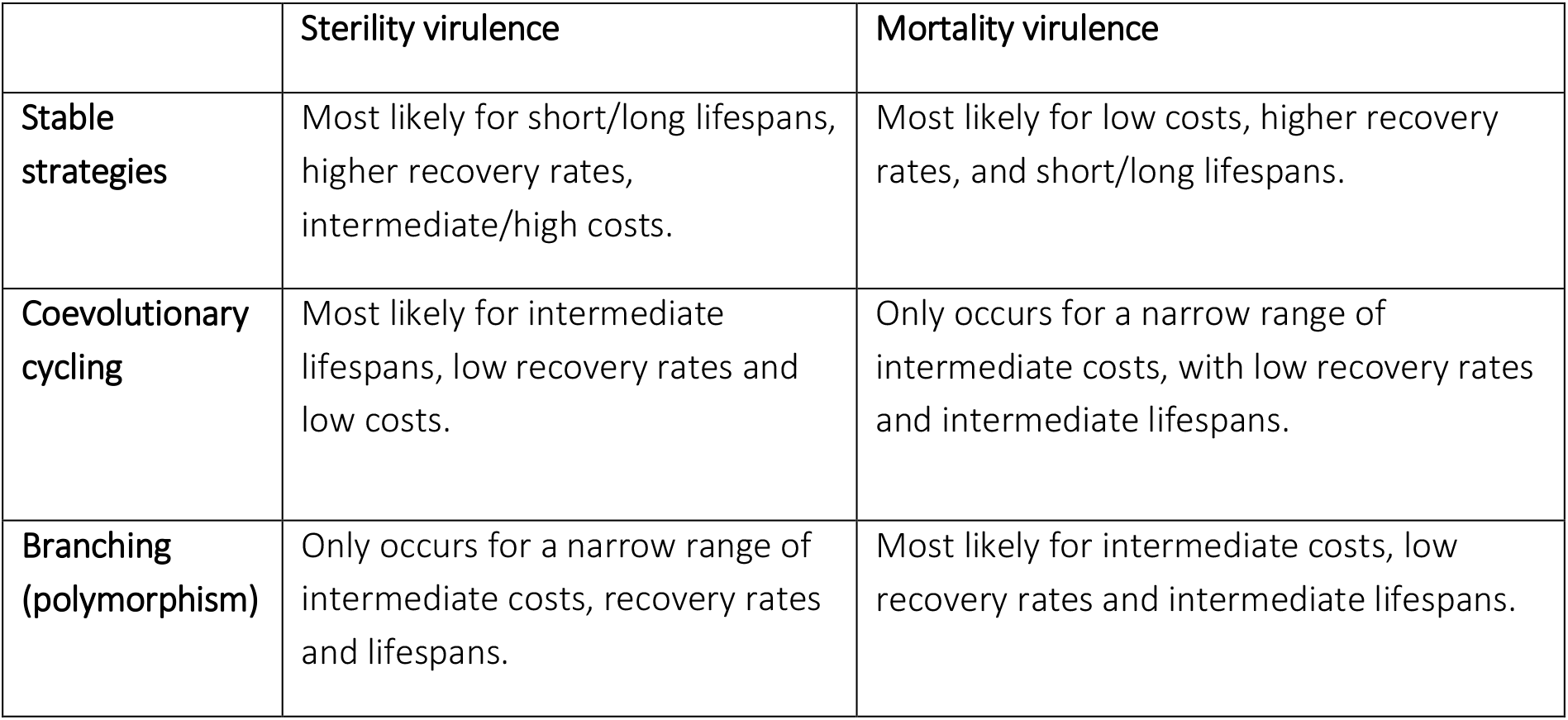
Summary of the conditions that favour the different coevolutionary outcomes.

|  | **Sterility virulence** | **Mortality virulence** |
| --- | --- | --- |
| **Stable strategies** | Most likely for short/long lifespans, higher recovery rates, intermediate/high costs. | Most likely for low costs, higher recovery rates, and short/long lifespans. |
| **Coevolutionary cycling** | Most likely for intermediate lifespans, low recovery rates and low costs. | Only occurs for a narrow range of intermediate costs, with low recovery rates and intermediate lifespans. |
| **Branching (polymorphism)** | Only occurs for a narrow range of intermediate costs, recovery rates and lifespans. | Most likely for intermediate costs, low recovery rates and intermediate lifespans. |

**Figure 4.**
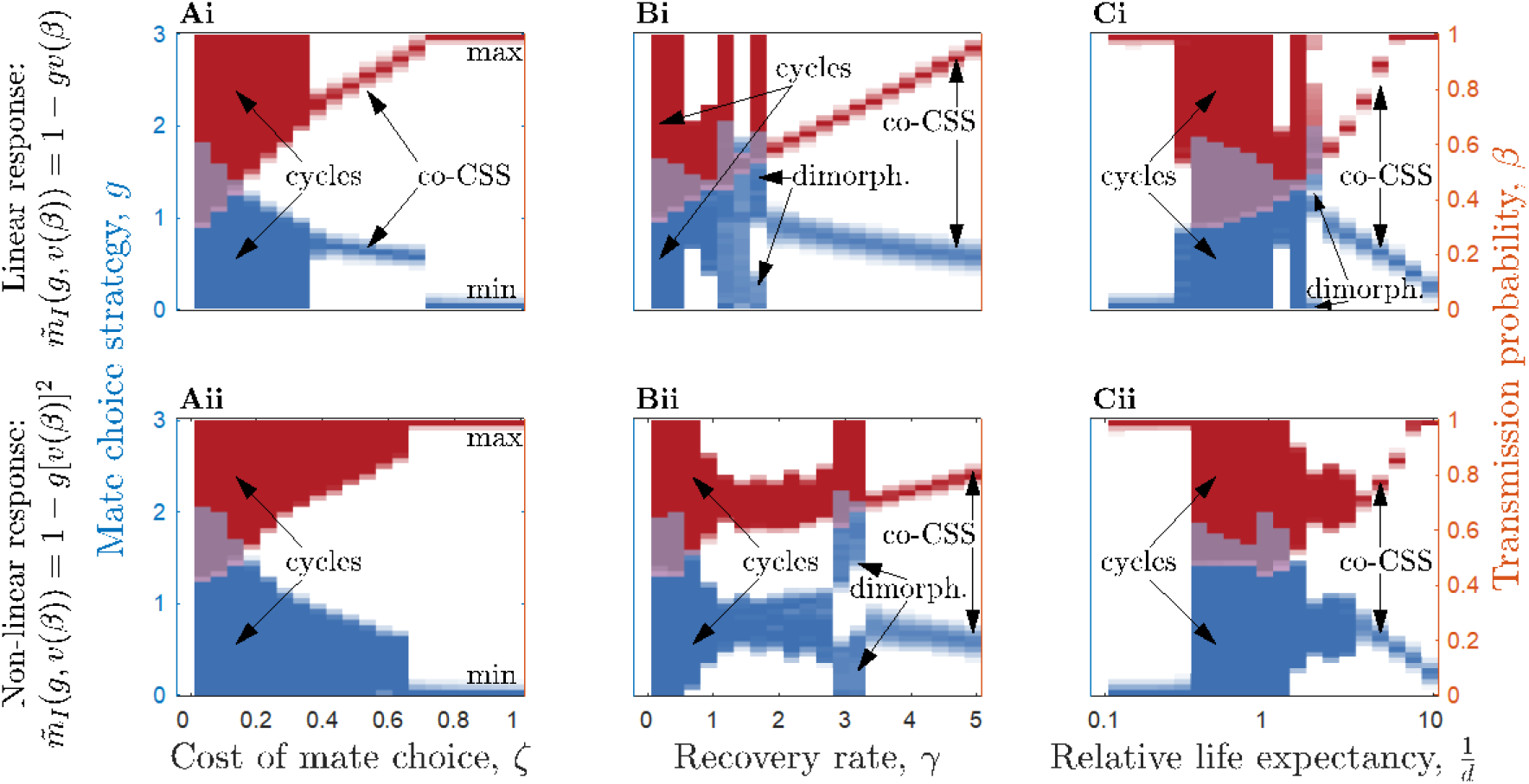
Qualitative and quantitative coevolutionary outcomes between the host mate choice strategy, ***g*** (blue), and the STI transmission probability, ***β*** (red), under sterility virulence (***𝑣***(***β***) **= 1** − ***f***(***β***) = ***βη***). Shading corresponds to the maximum frequency of each trait value over the final 2,000 evolutionary time steps (cycles are indicated by large solid regions). The top row corresponds to a linear mate choice function for 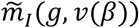 and the bottom row to a non-linear function of virulence. The outcomes are identified as: maximisation (max), minimisation (min), co-continuously stable strategies (co-CSS), coevolutionary cycling (cycles), and dimorphism (dimorph.). (Ai-ii) Effects of variation in the cost of mate choice, ***ζ***. (Bi-ii) Effects of variation in the recovery rate, ***γ***. (Ci-ii) Effects of variation in the relative life expectancy of the host, 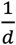. Fixed parameters as described in Fig. 3, with: (Ai-ii) ***γ*** **= 0.5**, (Bi-ii) ***ζ*** **= 0.1**, (Ci-ii) ***γ*** **= 0.5**, ***ζ*** **= 0.1**.

**Figure 5.**
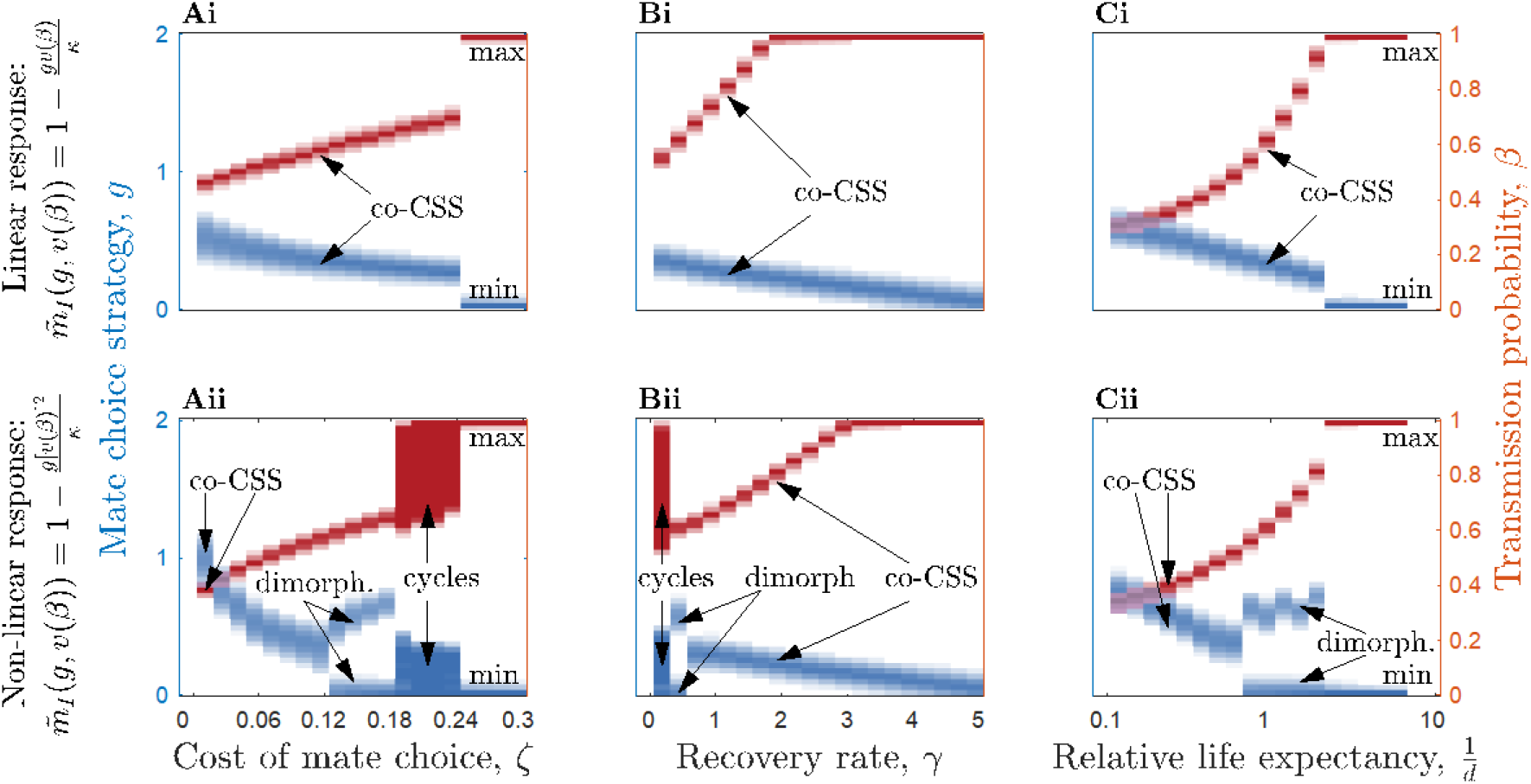
Qualitative and quantitative coevolutionary outcomes between the host mate choice strategy, ***g*** (blue), and the STI transmission probability, ***β*** (red), under mortality virulence (***𝑣***(***β***) **= 1** − ***α***(***β***) = ***βκ***). Shading and labels as described in Fig. 4. Fixed parameters as described in Fig. 3, with: (Ai-ii) ***γ*** **= 0.5**, (Bi-ii) ***ζ*** **= 0.15**, (Ci-ii) ***γ*** **= 0.5**, ***ζ*** **= 0.15**.

## Discussion

Understanding the role of STIs in the evolution of host mating strategies, and in turn, the effects of mating behaviour on disease evolution are inherently linked (Hamilton and Zuk 1982; Ashby and Boots 2015; Wardlaw and Agrawal 2019). Yet despite the large number of theoretical studies on coevolution between hosts and non-STIs, to date theoretical models of STIs have almost exclusively focused on one-sided adaptation rather than coevolution (Thrall et al. 1997, 2000; Knell 1999; Boots and Knell 2002; Kokko et al. 2002; Ashby and Gupta 2013; McLeod and Day 2014). In general there are more studies of evolution than coevolution, but there is a distinct lack of coevolutionary predictions for hosts and STIs compared to non-STIs (except see:Ashby and Boots 2015; Wardlaw and Agrawal 2019), which limits our ability to predict patterns of parasite-mediated sexual selection (PMSS).

Using a theoretical model of host-STI coevolution, I have shown that host mate choice can readily evolve under a broad range of conditions, including when hosts exhibit ephemeral mating dynamics, STIs cause either sterility or mortality virulence, hosts can recover from infection, and across large variations in host lifespan. In addition to showing when mate choice is most likely to evolve, I have also shown when qualitatively different coevolutionary outcomes usually occur (Table 1). Interestingly, coevolutionary cycling (fluctuating selection) in mate choice and STI virulence is much more common when the STI causes sterility virulence than mortality virulence, which may be because reductions in fecundity can cause sudden declines in population size and are generally known to induce oscillatory dynamics (Ashby and Gupta 2014). This suggests that STIs associated with higher mortality, such as dourine in equines (Gizaw et al. 2017), are more likely to lead to stable coevolutionary outcomes. But since STIs typically cause reductions in host fecundity (Lockhart et al. 1996) fluctuating selection may be a more probable outcome overall. Similarly, STIs often but not always cause chronic or long-lasting infections, which will promote coevolutionary cycling since lower recovery rates tend to have a destabilising effect. Higher clearance rates are associated with more stabilising outcomes, so for example we might expect acute STIs such as *Chlamydia trachomatis* in mammalian species (Miyairi et al. 2010) to produce more stable coevolutionary dynamics than STIs with low or zero clearance rates. Furthermore, fluctuating selection dynamics appear to be limited to hosts with moderate rather than short or long lifespans. Together, these predictions can help guide comparative analyses and other empirical studies towards systems with host and STI life-history traits that are more favourable to detecting these coevolutionary dynamics.

The model also revealed that evolutionary branching in host mate choice is possible, leading to the coexistence of more and less choosy individuals in the population. This only occurs when there are costs associated with being choosy, such that choosier individuals are not only less likely to mate with infected individuals but are also less likely to mate with susceptible individuals, for example due to false-positive detection of infection among healthy prospective partners. Hosts who are less choosy do not pay this cost but have a higher infection rate, which increases disease prevalence in the population. This is analogous to trade-offs with host life-history traits that are often associated with resistance or tolerance to infection (Schmid-Hempel 2003). Polymorphism in risky and prudent mating behaviour has been identified previously by Boots and Knell (2002), although their model only allowed for variation in the overall mating rate rather than for condition-dependent mate choice, and the mating rate for all infected individuals was the same regardless of whether hosts initially belonged to the risky or prudent mating type. While the model presented herein can also generate polymorphism (specifically, dimorphism), this time in terms of host mate choosiness (i.e. preferential mating for uninfected partners), polymorphism is predicted to be quite rare as it only occurs under a fairly narrow set of conditions, requiring intermediate costs, recovery rates, and host lifespans. This suggests that polymorphism in mate choice will be relatively rare in nature.

Previous theoretical work on host-STI coevolution appears to be limited to two studies. A recent paper by Wardlaw and Agrawal (2019) explored the coevolution of hosts and STIs in the context of sexual conflict, showing that the mode of virulence (sterility or mortality) can lead to contrasting coevolutionary outcomes, analogous to the present study. The other study, by Ashby and Boots (2015), is more closely related to the present model, as it too concerns the evolution of host mate choice for preferential mating with healthy hosts. Preferential mating with uninfected hosts is somewhat comparable to the notion of disease causing lower contact rates (e.g. due to decreased movement) in classical evolution of virulence theory (Ewald 1983), although there are also notable differences (sexual rather than direct transmission, and a reduction in contact rates affects reproduction). Ashby and Boots (2015) explored a pair formation model of host mate choice and sterility virulence in an STI, showing that transmission avoidance behaviour can evolve in the host and constrain STI sterility virulence, potentially leading to coevolutionary cycling in these traits. The present study differs from this work in two crucial aspects. First, the previous study focused on a system of serial monogamy with hosts moving between pools of paired and unpaired individuals, whereas the present study concerns the evolution of mate choice in hosts with ephemeral mating dynamics. All else being equal ephemeral mating dynamics likely represents the weakest selection for mate choice and therefore the strongest challenge to PMSS. This is because choosing a ‘bad’ mate (i.e. one who is infected, and potentially sterile) will have a less severe impact on lifetime reproductive success when sexual contacts are ephemeral compared to when individuals form a lasting contact, as the risk of infection will be higher than average and the reproduction rate lower than average while paired with an infected individual. Likewise, choosing a “good” mate (i.e. one who is not infected) will have a less significant positive effect on lifetime reproductive success when contacts are ephemeral, in part because individuals are no longer temporarily shielded from infection as they would be under serial monogamy. Despite the apparent reduction in the advantages of mate choice compared to serial monogamy, the results of the present study reveal it can still readily evolve under a wide range of conditions. To build on these results, future work should explore the evolution of mate choice under different mating systems, for example for polygynous or polyandrous hosts, as different mating systems have been shown to select for contrasting levels of virulence (Ashby and Gupta 2013). One especially interesting direction for future research that would unite the ephemeral and serial monogamy scenarios would be to examine the evolution of mate choice when serially-monogamous hosts engage in extra-pair copulations (Forstmeier et al. 2014). This would clarify whether mate choice should be stronger when choosing long- or short-term mating partners.

The second major difference with the model in Ashby and Boots (2015) is that here the STI can cause sterility or mortality and hosts may recover from infection, whereas the previous study only considered chronic STIs that cause sterility. Examining a broader set of STI characteristics helps to build up a more general picture of host-STI coevolution, which is important given that many STIs do not cause chronic infections or sterility virulence (Lockhart et al. 1996; Miyairi et al. 2010; Gizaw et al. 2017). As discussed above, the mode of virulence has an important impact on the nature of the coevolutionary dynamics, with mortality virulence tending to have a stabilising effect compared to sterility virulence. Overall, one might expect the benefits of mate choice to be greater if STIs cause sterility rather than mortality, as (1) individuals may be unable to reproduce again and (2) disease prevalence is likely to be higher as mortality virulence reduces ***R***_0_ by lowering the infectious period (equation 11), which means all else being equal the risk of infection will be lower under mortality virulence. Recovery from infection will typically reduce the benefits of mate choice as both disease prevalence and the costs of contracting an infection are lower (since infection is acute rather than chronic), yet recovery did not prevent the evolution of mate choice in the model. Instead, recovery tended to have a stabilising effect on the coevolutionary dynamics. As expected, high recovery rates reduce selection for mate choice, but surprisingly the difference in optimal levels of mate choice when infections are of relatively short or long durations is fairly small (Fig 4B, 5B), which supports the notion that both acute and chronic STIs can select for mate choice. For simplicity, I assumed that recovery does not lead to immunity from future infection and that the condition of recovered individuals does not differ from those who have yet to experience infection. The former assumption is reasonable for many STIs, which are less likely than non-STIs to result in lasting immunity (Lockhart et al. 1996), but the latter deserves further investigation. While it is possible for hosts to fully recover from infection, it is also reasonable to suspect that host condition may remain lower for some time following pathogen clearance, in which case these hosts should have lower mating success than individuals who have never been infected. In future, a simple extension of the current model would be to explore the effects of temporary or permanent reductions in host condition following infection, as this will separate the effects of mate choice into components representing transmission avoidance (i.e. avoiding infectious individuals) and partner fertility (i.e. choosing more fertile partners).

To date, many empirical studies have struggled to find evidence that hosts are able to discriminate between individuals with and without STIs (Abbot and Dill 2001; Webberley et al. 2002; Nahrung and Allen 2004). At first this seems surprising given that hosts should, in theory, be under strong selection to avoid choosing infected mates. There are a number of possible reasons as to why this may not always be the case. For example, hosts may simply be unable to detect signs of infection due to their own physiological limitations. This is not a particularly satisfying or general explanation, since various species have been found to prefer social or sexual contacts based on visual or olfactory cues relating to infection (Clayton 1990; Willis and Poulin 2000; Moshkin et al. 2012). Instead, it is more likely that hosts may be unable to detect infection due to strong selection on STIs to be inconspicuous, potentially through low virulence. For instance, sexually transmitted mites in ladybirds and the eucalypt beetle appear to have no negative impact on fertility or mortality under non-stress conditions (Webberley and Hurst 2002; Nahrung and Clarke 2007). Another related possibility is that hosts can sometimes discriminate between infected and uninfected individuals, but the costs of mate choice are too high relative to infection. In the model, mate choice only evolves under certain conditions, and may not evolve even when the STI is relatively virulent, conspicuous, or prevalent, if mate choice is intrinsically costly. Thus, any costs associated with mate choice (e.g. fewer mating opportunities) must be weighed against the potential benefits of avoiding infection. Another alternative is that hosts have other, more effective, forms of defence against STIs, such as post-copulatory grooming or urination to remove parasites (Hart et al. 1987; Nunn 2003). This area has received very little theoretical attention and is an intriguing target for future theoretical research on host-STI coevolution. Finally, it is possible that STIs cause changes in host characteristics such as attractiveness or behaviour such as mating frequency or choosiness, which may potentially counter or increase selection for mate choice. In principle such changes could simply be by-products of infection or could be host or STI adaptations to increase fitness by achieving a higher mating rate, potentially leading to sexual conflict (Knell and Webberley 2004; Apari et al. 2014; Wardlaw and Agrawal 2019). The evolutionary consequences for STIs increasing mating rates have yet to be thoroughly explored and deserve much greater attention in future theoretical work.

## Acknowledgements

I thank Sébastien Lion and Samuel Alizon for helpful comments on the manuscript.

## Notes

#### Summary of Updates

Rewritten and expanded throughout (introduction, methods, results, figures and discussion). Supporting information updated.

