## Supporting Information for "Antagonistic coevolution between hosts and sexually transmitted infections"

### DERIVATION OF HOST FITNESS

As described in the main text, the initial dynamics of a rare mutant host (subscript $m$) in a resident population at equilibrium are given by:

$$\frac{dS_m}{dt} = b_m - \beta[S_m I^*] - dS_m + \gamma I_m \tag{S1}$$

$$\frac{dI_m}{dt} = \beta[S_m I^*] - (d + \alpha(\beta) + \gamma)I_m \tag{S2}$$

with

$$b_m = \frac{p}{N^*}\big(r(1 - hN^*)(m_S(g_m)S^* + f(\beta)m_I(g_m, v(\beta))I^*)(m_S(g)S_m + f(\beta)m_I(g, v(\beta))I_m)\big) \tag{S3}$$

corresponding to the birth rate and

$$\beta[S_m I^*] = \frac{2\beta p m_S(g)m_I(g_m, v(\beta))S_m I^*}{N^*} \tag{S4}$$

to the infection rate.

One can use the next-generation method to calculate the invasion fitness for the rare mutant (Hurford et al. 2010), which first requires deriving the Jacobian of the mutant dynamics:

$$J = \begin{pmatrix} A - \frac{2\beta p m_S(g)m_I(g_m, v(\beta))I^*}{N^*} - d & B + \gamma \\ \frac{2\beta p m_S(g)m_I(g_m, v(\beta))I^*}{N^*} & -d - \alpha(\beta) - \gamma \end{pmatrix} \tag{S5}$$

where for notational convenience:

$$A = \frac{p m_S(g) r(1 - hN^*)(m_S(g_m)S^* + f(\beta)m_I(g_m, v(\beta))I^*)}{N^*} \tag{S6}$$

$$B = \frac{p m_I(g, v(\beta)) r(1 - hN^*)(m_S(g_m)S^* + f(\beta)m_I(g_m, v(\beta))I^*)}{N^*} \tag{S7}$$

Next, one must separate the Jacobian into components, $F$ and $V$ such that $J = F - V$ with:

$$F = \begin{pmatrix} A & B \\ 0 & 0 \end{pmatrix} \quad (S8)$$

$$V = \begin{pmatrix} \frac{2\beta p m_S(g) m_I(g_m, v(\beta)) I^*}{N^*} + d & -\gamma \\ -\frac{2\beta p m_S(g) m_I(g_m, v(\beta)) I^*}{N^*} & d + \alpha(\beta) + \gamma \end{pmatrix} \quad (S9)$$

The next-generation matrix is given by $N_G = FV^{-1}$, and the invasion fitness of the rare host mutant, $w_H(g_m, \beta)$, is sign equivalent to the largest eigenvalue of this matrix minus 1:

$$w_H(g_m) = \frac{prm_S^g(1 - hN^*)\left(2pf_\beta m_I^{g,\beta} m_I^{g_m,v_\beta} \beta I^* + \Gamma_\beta N^*\right)\left(m_S^{g_m} S^* + f_\beta m_I^{g_m,v_\beta} I^*\right)}{N^*\left(2p\beta m_S^g m_I^{g_m,v_\beta} I^*\left(\Gamma_\beta - \gamma\right) + d\Gamma_\beta N^*\right)} - 1 \quad (S10)$$

where $\Gamma_\beta = d + \alpha(\beta) + \gamma$, $f_\beta = f(\beta)$, $m_S^g = m_S(g)$ and $m_I^{g_m,v_\beta} = m_I(g, v(\beta))$, as shown in equation 31 in the main text.
